# In situ Structure of the Human Gap Junction

**DOI:** 10.1101/2025.07.04.663179

**Authors:** Evans Eshriew, Esa-Pekka Kumpula, Shiv K. Sah-Teli, Amiel Abettan, Amina Djurabekova, Vivek Sharma, Juha T. Huiskonen

## Abstract

Gap junction plaques (GJPs) enable direct intercellular communication and consist of connexin channels arranged into two-dimensional lattices. While structures of purified connexin channels have informed models of gating, they omit key intracellular regions and lack native context. Here, we use cryogenic electron tomography and focused ion beam milling to determine the in situ structure of human connexin-43 (Cx43) GJPs in HEK293 cells at 14 Å resolution. We discover a previously unresolved structural role for the large C-terminal domain in mediating lateral channel–channel interactions critical for plaque assembly. Coarse-grained molecular dynamics simulations reveal how lipids and cholesterol occupy the space between adjacent connexins. These findings resolve a decades-old question regarding gap junction organization and highlight a mechanistic function for the C-terminal domain, likely regulated by phosphorylation. Our study provides a structural blueprint for understanding how connexin diversity and regulation shape tissue-level communication in health and disease.

## Introduction

Gap junctions play a crucial role in intercellular communication by enabling metabolic and electrical coupling at cell–cell contact sites^1^. Gap junction plaques (GJPs) comprise hundreds of gap junction channels (GJCs) arranged into two-dimensional lattices that bridge the plasma membranes of adjacent cells^2^. These channels permit the bidirectional exchange of ions, secondary messengers, hormones, and other small molecules, and are involved in diverse processes such as cell differentiation, tumour progression and suppression, and cardiac conduction^3^. Each GJC is a dodecamer consisting of two hexameric connexin oligomers that dock head-to-head via their extracellular loops (**Figure 1a**)^4^. Of the 21 connexin isoforms identified in humans, each exhibits distinct conductance, permeability, oligomerization compatibility, regulatory mechanisms, and tissue distribution^5-7^. Mutations in connexin genes are linked to diseases including cataract, deafness, oculodentodigital dysplasia, keratitis-ichthyosis-deafness syndrome, Charcot–Marie–Tooth neuropathy and leukodystrophy^8^.

**Figure 1.**
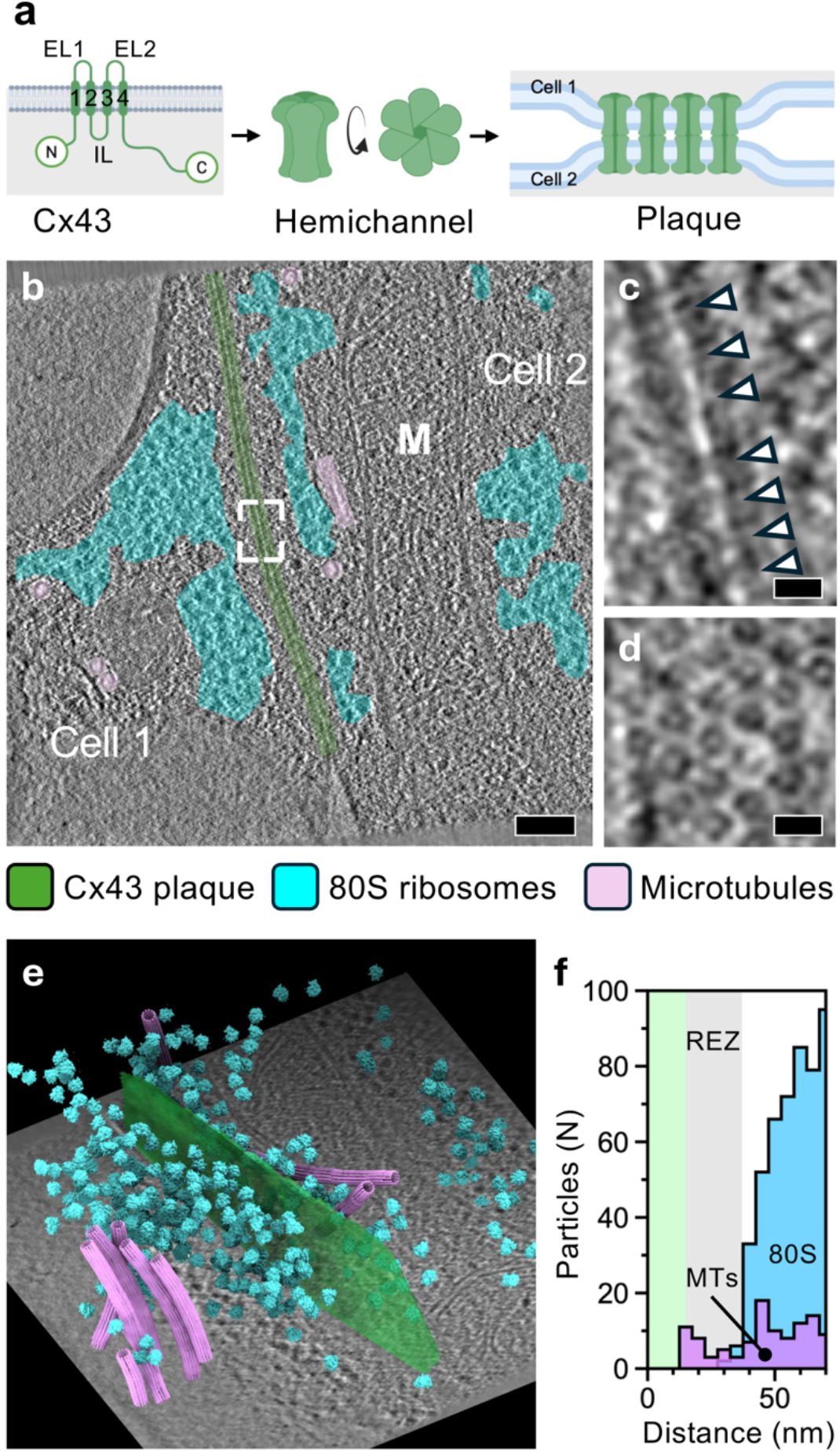
Cryogenic electron tomography and template matching of HEK293 Cx43-eGFP cells. (**a**) A schematic representation of the gap junction assembly from connexin 43 (Cx) monomers to hemichannels and to the gap junction plaque between two cells. Intra-(IL) and extra-cellular (EL) loops are labelled. The transmembrane helices are numbered 1–4. (**b**) A 1nm thick slice through a cryogenic electron tomography (cryo-ET) reconstruction of a cell-cell junction. Areas harbouring Cx43 channels (green), 80S ribosomes (cyan) and microtubules (pink) have been indicated. A mitochondrion has been labelled (M). Scalebar, 100 nm. (**c**) A close-up of the area indicated in *b*. Side views of Cx43 gap junction channels bridging two plasma membranes are indicated with arrowheads. Scalebar, 10 nm. (**d**) A close-up from a different cryo-ET reconstruction showing top views of the Cx43 gap junction channels. Scalebar, 10 nm. (**e**) The results of template matching for Cx43 gap junction channels, 80S ribosomes, and microtubule segments are shown after plotting the templates back onto the cryo-ET tomogram (illustrated with a 1nm thick slice in the background). The gap junction is depicted as a transparent surface (green) fitted to the Cx43 channel positions for visual clarity. (**f**) A histogram of pairwise distances between 80S ribosomes and microtubule segments (MTs) to the Cx43 gap junction channels, calculated from multiple cryo-ET reconstructions (N=26). The 15-nm-wide zone occupied by Cx43 channels is indicated in green. The 22-nm wide ribosome exclusion zone (REZ) is indicated in grey.

Gap junction plaques act as coordinated and dynamic platforms that enhance intercellular signalling. Their dynamics are influenced by a range of factors, including post-translational modifications, lipid composition, and interactions with regulatory proteins^9-16^. Moreover, GJPs, particularly those formed by connexin-43 (Cx43) have been shown to interface with cytoskeletal networks and cytoskeleton-associated proteins such as zonula occludens-1 (ZO-1) via the C-terminal domain (CTD)^17,18^. These interactions stabilize and modulate the size of GJPs^19^. Deletion of the Cx43 CTD results in the absence of GJPs highlighting its significance for assembly^20^. The impairment of these regulatory mechanisms has important implications for tissue physiology and disease. In cardiac arrhythmias, changes in GJP stability triggered by oxidative stress or altered connexin composition can disrupt electrical impulse conduction^21^. In cancer, dysregulation of GJPs has been linked to tumour progression^22^.

Current understanding of gap junction plaque structures has largely been informed by electron microscopy of immunogold-labelled freeze-fracture samples, immunocytochemistry, and studies using fluorescently tagged connexins, all of which have aimed to elucidate the structural organization, assembly mechanisms, and function of GJCs within plaques^23^. Additionally, atomic force microscopy has revealed conformational changes in purified connexin-26 plaques, particularly in the cytoplasmic domains and extracellular surfaces^24^. Despite these efforts, the molecular-level organization of GJPs and the specific interactions that drive their assembly remains poorly understood. More recently, cryogenic electron microscopy (cryo-EM) has provided a wealth of structural details on individual connexin channels shedding light on the effects of pathogenic mutations^25-27^, as well as pore dynamics^28,29^, pH-dependent gating^30^ and voltage-dependent gating^4^. However, this reductionist approach inherently omits the complexity of a native plaque environment. Notably, the CTDs have remained unresolved in these structures.

Here, we aimed to elucidate the structural architecture of the gap junction plaque in situ, focusing on Cx43 in human cells. To this end, we employed cryogenic electron tomography (cryo-ET) in combination with cryo-focused ion beam scanning electron microscopy (cryo-FIB/SEM) to visualize GJPs in their native cellular context. This approach enabled us to resolve molecular-level structural features of the GJC lattice in a near-native state. To gain further insight into the molecular interactions that underpin GJP formation and stability, we complemented our structural analysis with coarse grained (CG) molecular dynamics (MD) simulations. Our in situ structure of the connexin dodecamer in an open state, resolved at 14 Å resolution, revealed a key role for the C-terminal domains in mediating lateral channel–channel contacts and provides a blueprint for understanding the molecular mechanisms that govern gap junction plaque assembly.

## Results and Discussion

### Targeting gap junctions from micropatterned human cells for cryo-ET

To determine the ultrastructure of the gap junction plaque by cryo-ET, we established a workflow for preparing lamellae of cell–cell contacts by cryo-FIB/SEM^31^. We generated a HEK293T cell line stably expressing Cx43 tagged with enhanced green fluorescent protein (eGFP). Previously, it has been shown that C-terminally GFP-tagged connexins cluster normally to GJPs^32^. This tagging, however, has been shown to increase the GJP size in HeLa cells^33^. While advantageous for structural studies, this suggest that GFP-fusion may interfere with interactions to regulatory factors or the dynamics of GJP assembly or disassembly.

The HEK293T-Cx43-eGFP cells were grown on micropatterned cryo-EM grids (**SFigure 1**; see Methods). These grids enabled the enrichment of cells at the centre of the grid squares and prevented cell overcrowding, thereby improving the throughput of lamella preparation^34^. Of the 14 different patterns tested, an hourglass-shaped pattern proved advantageous for positioning GFP-positive cell–cell contacts in the middle of the grid squares. Cells on the micropatterned grids were subjected to fluorescence-targeted milling of 200-nm-thick lamella by cryo-FIB (**SFigure 2**). Correlating the GFP signal from cryogenic fluorescence light microscopy (cryo-FLM) with low magnification cryo-EM images confirmed the presence of cell–cell contacts in lamella. The GFP-positive cell–cell contacts were 3.4±1.4 μm (N=5) long. The GFP signal was interrupted by 0.4±0.2 μm (N=7) long gaps. A closer examination of the cryo-EM images revealed that in areas devoid of GFP signal, the cell–cell contact was absent, and the two membranes were dethatched and curved away from each other forming a cavity (**SFigure 2f**). These results show that HEK293T-Cx43-eGFP are suitable for targeting human GJPs by correlative imaging.

### Cryogenic electron tomography of the human gap junction plaque

Cryo-EM of 19 lamellae milled from 3 cryo-EM grids of adherent HEK293T-Cx43-eGFP cells revealed the presence of cubic ice, possibly due to the relatively large thickness of the cells in the areas of cell–cell contacts, resulting in incomplete vitrification (**SFigure 2g**). To overcome this for cryo-ET, we prepared grids in 10% glycerol as a cryo-protectant^35^. A total of 20 lamellae milled from such a grid showed no signs of cubic ice, suggesting that they were suitable for structure determination by sub-tomogram averaging. Interestingly, the continuous GFP-signals were shorter in the presence of 10% glycerol (0.5±0.4 μm; N=31) than in its absence (1.5±0.9 μm; N=13). A closer examination of the images revealed multiple punctate signals in the cytoplasmic regions. These signals likely correspond to connexosomes (annular gap junctions), vesicular structures formed by the endocytosis of GJPs^36^. Their internalization may have been induced by osmotic effects of 10% glycerol and left a vacuole-like discontinuity at the plasma membrane devoid of connexins and thus lacking GFP signal (**SFigure 2i**).

We used cryo-FLM for targeting 112 tilt series from the 20 lamella and reconstructed cryo-ET tomograms from 26 tilt series of suitable quality with a cell–cell contact region. The cell–cell contacts in these tomograms revealed repeated densities bridging the plasma membranes from the opposing cells (**Figure 1b,c**). In orthogonal view directions, these densities were seen to be ordered in a hexagonal lattice, reminiscent of the models of locally ordered connexin lattices (**Figure 1d**)^37^. As these structures are positive for the GFP signal, we attribute them to GJPs. Visual inspection of the tomograms revealed, in addition, organelles in proximity to the GJPs, such as a mitochondrion (**Figure 1b**) and a Golgi apparatus (**SFigure 4a**). Furthermore, vesicles, possibly involved in trafficking, were seen in proximity to microtubules and GJPs (**SFigure 4b**,**c**). In one instance, a coated vesicle, possibly in the process of membrane fusion, was connected to the plasma membrane next to a GJP (**SFigure 4d**). We also observed a connexosome (400 nm in diameter) in the process of being internalized by endocytosis (**SFigure 4e**,**f**). These observations suggest that the HEK293T-Cx43-eGFP cells supplemented with 10% glycerol are a suitable model system for studying the cellular architecture in the context of human GJPs.

To map the molecular arrangement of the gap junctions, we used template matching to detect the positions of Cx43 GJCs, in addition to 80S ribosomes and microtubules, in 26 tomograms (**Figure 1e**). To account for possible effects of 10% glycerol on structure determination from these samples, as a control, we determined the structure of the 80S ribosome to 10 Å resolution from 6135 particles (**SFigure 3**). This and an earlier 4.9-Å average of 80S ribosomes calculated using a significantly larger data set (15,628 particles) from HEK293 cells with 10% glycerol^35^ show that samples prepared this way are suitable for sub-tomogram averaging. We further quantified the distances of ribosomes, in addition to microtubules, to the GJCs by calculating their closest pair-wise Euclidean distances (**Figure 1f**). Ribosomes were apparently excluded from the vicinity of the GJPs within a 22-nm-wide ribosome exclusion zone (REZ). This suggests that the relatively large Cx43 C-terminal domains, and possibly other proteins interacting with this domain, in addition to actin filaments^13^, occupy this region. Interestingly, microtubules were occasionally present within REZ (**Figure 1f**), consistent with their reported interaction with Cx43^17,38^.

### Curvature and asymmetry in the gap junction channel lattice

To determine the ultrastructure of the gap junction in situ, we used template matching followed by sub-tomogram averaging. Classification of 30,338 sub-tomograms of GJC lattice patches from 26 cryo-ET tomograms revealed lattices with varying degrees cylindrical and saddle curvatures (**SFigure 5a–c**). The direction of the connexin lattice was observed to be uncoupled from the principal axes of curvature. Conversely, connexin patches with the same curvature and lattice orientation clustered in the same local areas of the GJP (**SFigure 5d–f**). We extracted, aligned, and averaged 11,904 sub-tomograms with a clear GJC channel lattice to reconstruct a local patch of the gap junction channels at 14-Å resolution (**Figure 2, SFigure 3**). The volume, reconstructed with C6 symmetry, reveals a lattice of Cx43 channels (**Figure 2a– c**). Each channel is formed by two opposing Cx43 hexamers as has been reported earlier (**Figure 2d**)^39^. We quantified the density distribution in the model, considering the radius of curvature (1.05 µm) of the GJC lattice in this reconstruction. An additional layer of density was observed on the concave side, but not the convex side, of the GJC lattice possibly corresponding to other proteins inducing or sensing the negative curvature (**Figure 2d–e**). The total thickness of the Cx43 GJP measures ∼30 nm. The channel-to-channel distance, measured from the channel centres, is 90 Å, consistent with a previous report (**Figure 2f**)^40^.

**Figure 2.**
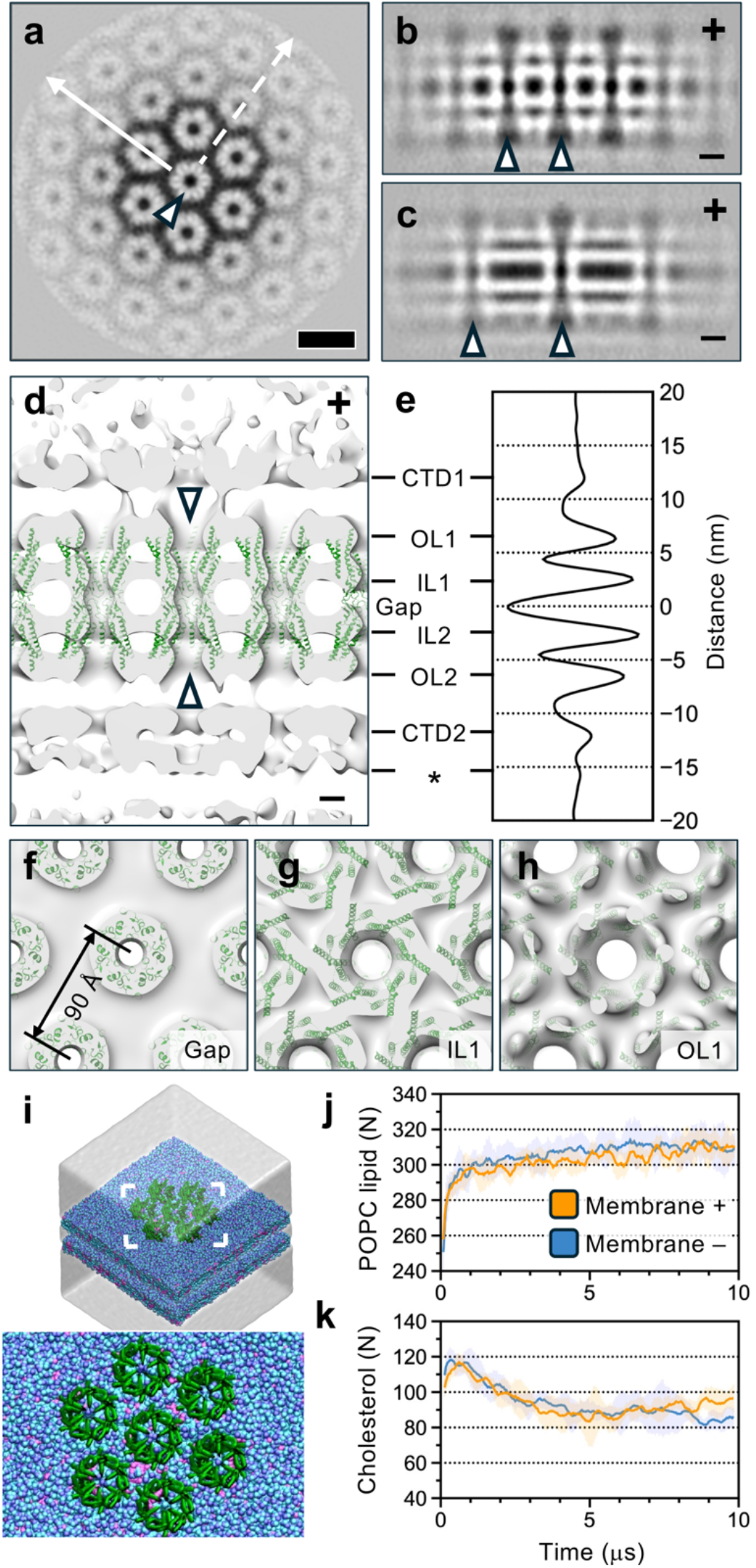
Sub-tomogram averaging of Cx43 gap junction channels. (**a**) A 12-Å -thick slice through a sub-tomogram average of the Cx43 channel lattice is shown in the plane of the membranes. One channel is indicated with an arrowhead. Scalebar, 10 nm for *a*–*c*. (**b**) A 12-Å-thick slice through the sub-tomogram average, along the solid line in *a*, is shown from the side. The convex (+) and concave (–) sides of the gap junction patch are indicated. (**c**) A 12-Å-thick slice, along the dashed line in *a*, is shown. Two channels are indicated with arrowheads in both panels. (**d**) An isosurface rendering of a sub-tomogram low-pass filtered to 20-Å resolution is shown as a grey transparent surface. The fitted structures of Cx43 channel atomic model (PDB:7Z22, residues 17–105 and 151–235) are shown in green ribbon. Two hemichannels forming a full channel are indicated with arrowheads. The convex (+) and concave (–) sides of the gap junction patch are indicated. (**e**) A density distribution as a function of distance from the gap junction mid-point is shown. Density is in arbitrary units. The inner (IL) and outer (OL) leaflets of the two lipid bilayers are marked. The regions corresponding to the C-terminal domains (CTD) of Cx43 hexamers are labelled. An additional layer of density on the concave side of the gap junction is indicated with an asterisk. (**f–h**) Isosurface renderings are shown from the top (+ to – direction) for the extracellular region (gap), inner leaflet (IL1) and intracellular densities (CTD1). Three of the six putative lateral contacts between individual channels at the local two-fold axes of symmetry are indicated with arrows. (**i**) A rendering of the coarse-grained molecular dynamics simulation setup. The connexins are green, the POPC lipids are blue (head groups in a darker shade), cholesterol is magenta. The solvent water is rendered as a transparent cube. The inset shows the area indicated. (**j–k**) The number of POPC lipids (*j*) and cholesterol molecules (*k*) around the centremost gap junction channel is plotted as a function of the simulation time for both the membranes (+ and – sides).

### Intracellular ordered structural elements bridge gap junction channels

To understand channel–channel interactions in the lattice, we fitted an atomic model of the Cx43 channel^25^ into the density, with a cross-correlation coefficient of 0.59 both for the central dodecamer and its neighbour, which were fitted independently. This fitting established both the positions and the in-plane orientations of the GJCs in the lattice **(Figure 2d**). Importantly, fitting of this GJC model, which encompasses the transmembrane regions in addition to the extracellular loops (covering approximately 50% of the entire structure; see **Figure 1a**), revealed no contacts between individual GJCs (**Figure 2f–h**). Instead, coarse grained molecular dynamics simulations showed that on average approximately 16 cholesterol and 50 POPC molecules reside between any two connexin hemichannels (**Figure 2g–i**). A corollary of this finding is that the lateral channel–channel contacts needed to assemble an ordered Cx43 GJC lattice must reside in the intracellular region encompassing the intracellular loop (IL; residues 105–150) and the C-terminal domain (CTD; residues 236–382). Consistent with this notion, ordered density resides proximal to both membranes (**Figure 2d**). We assign this density to the Cx43 CTDs and the ILs. We cannot, however, exclude the possibility that the C-terminal eGFP-tag or connexin-interacting proteins also contribute to this density. Notably, the CTD is disordered in the Cx43 GJC determined using purified channels^25^. Our data suggests that it becomes at least partially ordered in the cellular context and that it interacts with additional cellular factors, especially on the concave side of the slightly curved membrane.

Our Cx43 GJC lattice model facilitates the comparison to other modelled connexin lattices (**SFigure 6**). Cx26 is one of the smallest connexins (226 residues) and largely lacks the CTD. It has been reported to have much closer channel-to-channel distances (77±5 Å)^24^ compared to Cx43 (90 Å). Cx26 GJC lattice model suggest that the channel–channel contacts are created by the short C-terminal tail (222–227) possibly in addition to the transmembrane helix 4 (218– 221). This difference suggests that Cx43 has evolved a looser lattice where contacts are mediated by its extended CTDs, possibly to allow for a finer regulation of the GJP.

### In situ structure of the human gap junction channel

The in situ structure of the gap junction channel shows a varying degree of order (**Figure 3**). The extracellular loops connecting the two hemichannels, in addition to the transmembrane (TM) helices 1 and 2, are relatively well ordered. In contrast, TM helices 3 and 4 are disordered, suggesting relatively higher mobility, structural variation or deviation from a strict six-fold symmetry in these helices that are distal from the channel pore (**Figure 3a–c**). To determine whether the pore observed in situ is open or closed, we compared its cryo-ET map to the previously published single-particle cryo-EM structure of Cx43 GJCs in nanodiscs (**Figure 3d**)^25^. We first low-pass filtered this map to 14 Å resolution to facilitate comparison to our cryo-ET map at this resolution. In this structure, the N-terminal helices (residues 1–16) form a lid that closes the channel, and this feature is resolved at 14 Å resolution. As this lid feature is absent in our in situ structure, we conclude that the channels are predominantly in an open conformation in our in situ data. Due to averaging approach applied, we cannot, however, exclude the possibility that a fraction of the channels is in a closed or an intermediate conformation.

**Figure 3.**
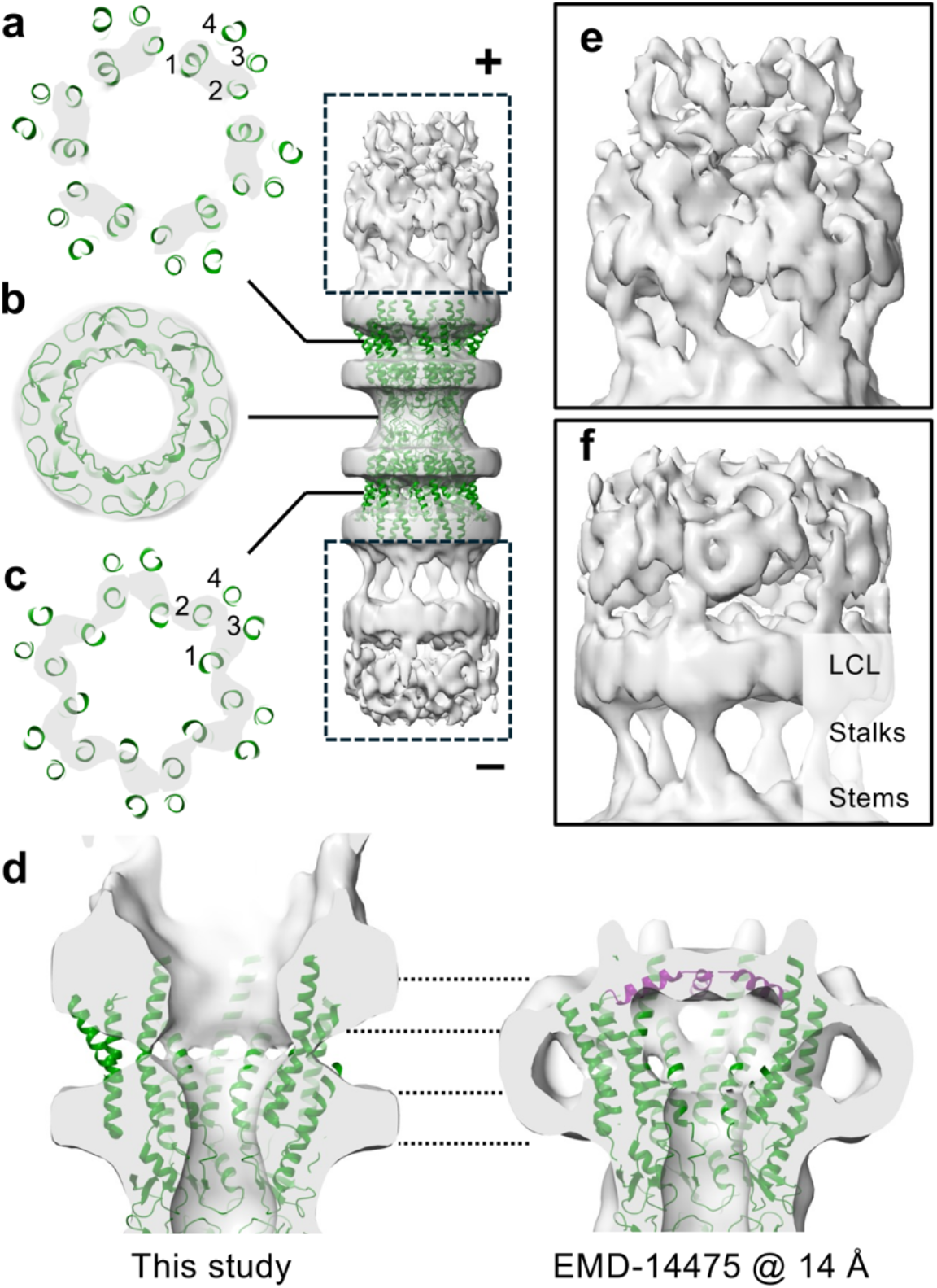
Structure of the Cx43 channel. (**a–c**). An isosurface rendering of the cryo-ET sub-tomogram average at 14 Å resolution is shown as a grey transparent surface, together with a partial atomic model of the Cx43 channel (PDB:7Z22, residues 17–105 and 151– 235). The inset indicates the level of the cross sections taken from the middle of the top bilayer (*a*), the middle of the extracellular loops (*b*) and the middle of the bottom bilayer (*c*). The transmembrane helices are numbered 1–4 for one Cx monomer in both hemichannels. The convex (+) and concave (–) sides of the gap junction are indicated. (**d–e**) Close-ups of the intracellular areas indicated in the inset (dashed rectangles) are shown. In both, a stalk-like density can be seen, connecting the density below it (stem) and the density above it (LCL, lateral contact layer). (**f–g**) Close-up of the hemichannel cryo-ET map (*f*) and a single particle cryo-EM structure of the Cx43 hemichannel, low-pass filtered to the same resolution (14 Å) for comparison. At this resolution, the N-terminal helices (residues 1–16, purple) closing the channel are visible in the cryo-EM density. The dashed lines indicate the position of the two membrane leaflets in both structures.

Both intracellular regions proximal to the membranes show partially ordered density (**Figure 3e–f**). In both regions, a stalk-like density is seen protruding from a relatively bulkier region termed “stem”, and connecting to a layer of density termed “lateral contacts layer” (LCL). This layer of density extends over neighbouring channels, most likely creating the channel–channel contacts. This layer most likely includes the CTDs due to their extensive size (residues 236–382). Fitting of a predicted full-length Cx43 GJC structure places the IL in the stem region of the intracellular density (**SFigure 7**). Furthermore, a putative helix-loop-helix motif (HLH, R299–E352) is predicted within the CTD (**SFigure 7**). Nuclear magnetic resonance (NMR) experiments of the CTDs have shown that in human Cx43 S314–I327 and Q342–A348 exhibit α-helical structure^41^. Furthermore, a HLH motif (A315–A348) has been detected in rat Cx43 CTD^42^, which has a high degree of sequence identity to that of human Cx43 (95%). We hypothesized that this motif may create either dimeric or trimeric channel– channel contacts. Consistent with this hypothesis, Cx43 C-terminal deletion mutant α236–382 fails to assemble gap junction plaques^20^. Furthermore, NMR experiments have shown that three regions within this motif (R299–Q304, S314–I327, and Q342–A348), in addition to one preceding it (M281–N295), have the tendency to drive Cx43 CTD dimerization at low pH^43^. We propose that the residues before the HLH (236–314) contribute to the relatively ordered stem and stalk regions observed in our cryo-ET structure. Finally, the superposition of the predicted Cx43 GJC full-length model on our GJC lattice revealed a putative network of lateral interactions between HLHs motifs (**SFigure 7f**). Taken together, these observations suggest that the Cx43 CTD creates the channel–channel interactions required for the GJP assembly via the dimerization of the putative HLH-motifs.

## Conclusions

By establishing a cryo-ET workflow in HEK293T cells expressing connexin-43, we enabled direct visualization of connexin lattices within human gap junction plaques in situ. Cryoprotection with glycerol preserved the ultrastructure of cell–cell contacts and allowed high-resolution imaging of gap junction plaques. These plaques adopt a semi-ordered lattice of gap junction channels embedded in a variably curved membrane. The lattice exhibits further structural asymmetry marked by additional density on the concave cytoplasmic face, potentially corresponding to binding partners or curvature-sensing components. As individual connexin-43 channels are not in direct contact, but instead spaced by lipids and cholesterol, we propose that lateral channel–channel contacts are mediated by C-terminal domains. These domains, disordered or absent in previous studies of detergent-purified channels, are ordered in situ and appear to be essential for lattice gap junction assembly. Higher resolution will be required to resolve their conformations and molecular contacts in detail. Furthermore, the physiological significance of the HLH motif needs to be established in further studies. In conclusion, our study provides a framework for understanding the spatial organization of gap junctions in the cellular context.

## Methods

### Connexin 43 cloning and stable cell line generation in HEK293T

Human connexin 43 (Uniprot ID P17302) plasmid was obtained from the Human ORFeome Library (Genome Biology Unit, Helsinki Institute of Life Science, University of Helsinki). Connexin 43 with a C-terminal 3C-eGFP-SpyTag003^44^ tag was cloned into pURD vector^45^ and used for stable cell line generation in HEK293T cells. HEK293T cells were cultured in Dulbecco’s Eagle Modified Medium (DMEM) supplemented with 10% fetal bovine serum (FBS, Gibco), 5% non-essential amino acids (NEAA, Gibco), 5% L-glutamine (Gibco) and incubated at 37°C in a 5% CO_2_-containing atmosphere. To obtain HEK293T cells stably expressing Cx43-3C-eGFP, cells were transfected with 1 µg of pURD vector harbouring Cx43-3C-eGFP and 3 µg of integrase expression vector (pgk-φC31/pCB92) using Lipofectamine 2000 (Thermo Fisher Scientific). Confluent cells were selected using 4 µg/mL puromycin (Thermo Fisher Scientific) for 2 weeks. For clonal line generation, GFP-positive cells were isolated by fluorescence-activated cell sorter (FACS) instrument (FACSAria II, BD Biosciences). Clones were monitored, expanded, and the fluorescence intensity was validated using a plate reader.

### Micropatterning of cryo-EM grids

Cryo-EM grids (Quantifoil R2/2 Au 200 mesh with a SiO_2_ foil) were micropatterned using a micropatterning system (Alvéole PRIMO). The grids were plasma cleaned at 0.8 Torr for 50 seconds using a tabletop high-power expanded plasma cleaner (Harrick Plasma PDC-002). For micropatterning the grids, a two-step passivation approach was used. First, the grids were transferred onto a glass coverslip covered with parafilm, and 20 μL of 0.01% of poly-L-lysine solution (CAS number 25988-63-0, P4707, Sigma-Aldrich) was immediately added. The glass coverslip containing the grids was transferred into a humidified chamber and incubated overnight at 4°C. Next, the grids were washed three times in 0.1 M HEPES pH 8.5 and incubated in 20 μL of mPEG-succinimidyl valerate solution (50 mg/mL in 0.1 M HEPES pH 8.5, MW 5000, mPEG-SVA, Laysan Bio) for 1 hour at room temperature. The grids were washed three times in deionized water. Excess water from each grid was removed by blotting and the grids were air dried. A 3-μL aliquot of a 1:3 dilution solution of photoinitiator (PLPP gel, Nanoscalelabs) in 70% ethanol was added to the surface of each grid and incubated in the dark until the gel was completely dry. The grids, fixed in position on a glass coverslip by PDMS stencil with the SiO_2_ foil side up were placed on an inverted widefield microscope (Nikon Eclipse Ti-E) mounted with Alvéole PRIMO. Each grid was then exposed to UV light at 100mJ/mm2. After micropatterning, the grids were washed three times in deionized water and further washed three times in 1×PBS. The grids were kept in a 35-mm dish, hydrated in 1×PBS, and stored at 4°C until they were ready to be used.

### HEK293T cell culture and seeding

HEK293T cells stably expressing Cx43-3C-eGFP were maintained in DMEM with 10% FBS, 5% NEAA, 5% L-glutamine and incubated at 37°C in a 5% CO_2_-containing atmosphere. The micropatterned grids were sterilized in 70% ethanol for 5 min under UV light and subsequently washed three times in deionized water and three times in 1×PBS. To functionalize the micropatterned grids, 20 µL ECM solution containing 15 µL fibronectin (cat number 1918-FN-02M, Bio-Techne) mixed with 5 µL fibrinogen-647 (fibrinogen from human plasma, Alexa Fluor-647 conjugate, Life Technologies Corporation) at 3:1 ratio in 1×PBS was added to each grid and incubated at room temperature for 1 hour. The ECM solution was removed from the grids by using pipette and the grids were washed three times in 1×PBS. The grids were conditioned in DMEM by washing three times. HEK29T cells were trypsinized, resuspended in DMEM, and filtered through a 40-µm cell strainer. A 5-µL aliquot of cell suspension at a concentration of 5×10^4^ cells/mL was added to each grid in drops sequentially until a minimum of 2–3 cells per grid square was achieved. After 2 hours, grids were washed with DMEM and incubated at 37°C in 5% CO_2_-containing atmosphere for approximately 24 hours before vitrification.

### Cryo-FIB/SEM lamella preparation

The cells were vitrified into liquid ethane using a manual plunge freezer. Optionally, 5-µL of 10% glycerol diluted in 1xPBS was added to the cells immediately before plunge freezing. Cryo-FIB of frozen hydrated HEK293T cells was performed using an Aquilos2 dual-beam FIB-SEM microscope (Thermo Fisher Scientific). Metallic platinum was deposited on the grids (1 kV, 10 mA, 10 Pa, 15–20 seconds) followed by organic platinum deposition via a gas injection system for 10–20 seconds. AutoTEM-cryo software (Thermo Fisher Scientific) was used to mill the cells with a milling angle of 12° to a thickness of 1 µm in three steps with decreasing ion beam currents (1, 0.5, and 0.3 nA) at 30 keV. Lamellae polishing was achieved by thinning to a final thickness of 200–300 nm at 50 pA. To complete the milling process, the lamellae were sputter-coated with platinum (1 kV, 10 mA, 10 Pa, 5–15 s). To prevent the lamella from breaking or bending, microexpansion joints were used during milling.

### Cryo-fluorescence light microscopy imaging of the lamellae

The milled EM grids were imaged using a super-resolution confocal microscope (Zeiss LSM 900 Airyscan) equipped with a cryo-stage (Linkam). A low-magnification air objective (5×) in wide-field mode was used to acquire an overview image of the lamellae. This was followed by acquiring z-stacks with a 0.5-μm spacing covering 4–6 μm in Airyscan mode with 488 laser line and a 100×/0.75 NA air objective (Zeiss Plan-Neofluar), using a pixel size of 79 nm. Reflection-mode images were acquired for each z-plane. To increase the signal-to-noise ratio of each image stack, the stacks were averaged, and maximum intensity projections were generated. Correlation of acquired Airyscan images and low-magnification TEM images (lamella maps) was performed with Icy eC-CLEM^46^ and Fiji BigWarp^47^ plugins by using the lamellae shape and features visible in the reflection images as landmarks.

### Cryo-ET data acquisition

Cryo-ET data were collected on a Titan Krios G4 transmission electron microscope operated at 300 kV and equipped with a Falcon4i direct electron detector and a SelectrisX energy filter (Thermo Fisher Scientific). The lamella pretilt axis was aligned with the microscope stage tilt axis. Data were acquired using SerialEM software (v 4.1.0beta)^48^ and PACE-tomo^49^. To record 112 tilt series from 20 lamella, a dose-symmetric tilting scheme^50^ from –40° to +64° with 2° increments was used, starting from the lamella pre-tilt angle (12°). Tilt series were acquired in counting mode at 3–6 μm under focus using a pixel size of 2.4 Å/pixel. For each tilt, an electron exposure of 3 e^−^/Å^2^ per tilt was used, resulting in a total electron exposure of approximately 160 e^−^/Å^2^.

### Tomogram reconstruction and sub-tomogram averaging

Cryo-ET data were processed using the Warp-RELION-M pipeline^51^. A subset of 26 tilt series was selected from the total of 112 by visual inspection for further processing. After motion correction and contrast transfer function (CTF) estimation of the frame series in WarpTools (v2.0.0), tilt images were filtered by defocus and CTF resolution, retaining images with a defocus higher than 2.6 μm and resolution of 11 Å or better. Down-sampled tomograms (9.6 Å/pixel) were reconstructed for template matching in PyTom Python package pytom-match-pick^52^. An initial template volume of the connexin lattice was generated by tracing the gap junction regions manually in Dynamo (v.1.1.532)^53^ and averaging the patches with D6 symmetry in RELION (v5.0beta)^54^. For template matching of the 80S ribosome and microtubules, EMD-15815 and EMD-6351 were used, respectively. The templates were low-pass filtered to 40-Å resolution. The coordinates were extracted using a top-hat filter^52^. The coordinates for connexin patches and microtubule segments were manually curated in UCSF ChimeraX^55^ using ArtiaX plugin^56^. For connexin patches, only the coordinates corresponding to a 2D surface matching the gap junction region in the tomograms were retained. For microtubule segments, only the coordinates corresponding to 1D tracks of the filaments were retained. For connexins, the tilt and psi Euler angles were added as priors using a custom Python script. The connexin and ribosome particles were exported to RELION using Warp at 4.8 Å/pixel sampling and a box size of 128 pixels to classify and refine the particles. For connexins, C2 symmetry was applied to retain different curvatures of the connexin patch. The resulting averages were used for the second round of template matching in pytom-match-pick and manual curation in ArtiaX. Further classification in RELION revealed two types of curvature in the connexin patches. These two templates were used for the third and final round of template matching in PyTom. The coordinates were consolidated from two separate template matching runs, keeping the highest correlating match for each position using a custom Python script. After refinement of all data sets in RELION, the connexin, ribosome and microtubule particles were exported to M for multi-particle refinement^51^. The particles were refined in seven stages: 1. Five rounds of 1×1 image warp with particle refinement. 2. Five rounds of 4×4 image warp with particle refinement. 3. Per-tomo weighting followed by five rounds of particle refinement. 4. Per-tomo weighting followed by five rounds of particle refinement. 5. Exhaustive defocus parameter estimation followed by five rounds of 4×4 image warp with particle refinement. 6. Per-tomo weighting followed by five rounds of particle refinement. 7. Five rounds of 4×4×1×10 volume warp with particle refinement. The refined connexins and ribosome particles were exported to RELION at 2.4 Å/pixel sampling and using a box size of 256 pixels for a final round of refinement. C6 symmetry was imposed on the connexin map. The Fourier shell correlation (threshold 0.143) was used to estimate both the global and local resolutions from two independent half datasets in RELION. The connexin map was sharpened using an ad hoc inverse B-factor of –500 Å^2^. The cryo-EM maps, including that of the microtubule segment at 40-Å resolution, were plotted back on the tomograms in ChimeraX using ArtiaX plugin. The ribosome-to-connexin and microtubule-to-connexin distances were calculated using a custom Python script. The density distribution of the gap junction was calculated from a cryo-EM reconstruction low-pass filtered to 20 Å resolution within an aperture of 15 nm and considering the radius of curvature 1.05 µm using a custom Python script. The radius and directions of curvature were determined by fitting a surface (a sphere, a cylinder or a saddle) to the centres-of-mass of Cx43 models rigid body fitted to the 3D class averages (with C2 symmetry) or the final connexin map (with C6 symmetry) using a custom Python script.

### Molecular modelling and coarse-grained molecular dynamics simulations

To probe the dynamical behaviour of lipids and their interactions with Cx43 GJCs, we created a coarse-grained model based on the 14 Å cryo-ET structure of Cx43 GJC lattice with *martinize2* software (version 2.9) and using the Martini 3 forcefield^57^. The residues corresponding to the intra-cellular loop (106–150) were modelled with AlphaFold2^58^. The double membrane CG model of the protein complex comprises seven Cx43 hemichannels immersed in each of the two POPC/cholesterol containing phospholipid bilayers. The phospholipid bilayer consisted of 70% and 30% POPC and cholesterol, respectively. The CG protein–membrane system was solvated with the standard Martini 3 water and ion beads with a 150 mM concentration of Na^+^ and Cl^−^ ions using the *insane*.*py* script^59^.

All MD simulations were carried out using the GROMACS software (version 2022)^60^. To avoid any steric clashes, the CG system was energy minimized for 5000 steps. The arrangement of the GJCs was restrained in production simulations by applying a harmonic force constant of 1000 kJ mol^-1^ nm^-2^ on the backbone CG beads. The velocity-rescaling thermostat^61^ and c-rescaling barostat^62^ were used to maintain the temperature at 300 K and pressure of 1 bar, respectively. The reaction field algorithm was used to treat the coulombic interactions using *ε*_*r*_ = 15^63^. Implementation of the Verlet cut-off scheme was utilized with a Lennard Jones cut-off of 1.1 nm^64^. Three 10 µs CG MD simulations were performed using a timestep of 20 fs.

To estimate the number of lipids located between adjacent connexin hemichannels, we performed analysis using MDAnalysis^65^. First, the center-of-mass (COM) of the central hemichannel was selected, and the distance to the COM of its adjacent hemichannels in the same membrane was used to define a radial cutoff (8.5 nm). All lipids (either POPC or cholesterol) within this radial region were selected and counted along three simulation trajectories of 10 µs each. The total number of lipids within the defined cutoff was then divided by six to approximate the number of lipids, i.e. POPC and cholesterol molecules shared between each pair of adjacent hemichannels. This procedure was applied independently for each membrane bilayer.

We predicted structure of the full-length structures of Cx26 and Cx43 with D-I-Tasser^66^. Dodecamers were created by matchmaker command in ChimeraX, using the atomic coordinates PDB:7QEQ and PDB:7Z22, respectively, as templates. Phosphorylation sites were retrieved from PhosphoSitePlus^67^ and mapped onto the structure in UCSF ChimeraX.

## End notes

## Acknowledgements

We thank Maryna Green, Xun Lu, Kiran Ahmad, Tuomas Niemi-Aro, and Zhengyi Yang for technical assistance. This study was funded by grants from the Sigrid Jusélius Foundation (to J.T.H, to V.S.), the Jane and Aatos Erkko Foundation (to V.S), the Research Council of Finland (to V.S) and the Finnish Society of Sciences and Letters (to V.S). This work benefited from access to the Electron Microscopy and Cryo-CLEM, Heidelberg, EMBL, an Instruct-ERIC centre. Financial support was provided by Instruct-ERIC (PID 25466). The flow cytometry analysis was performed at the HiLIFE Flow Cytometry Unit, University of Helsinki. We acknowledge CSC – IT Center for Science, Finland, for generous computational resources. We acknowledge the facilities and expertise of the HiLIFE CryoEM unit at the University of Helsinki, a member of Instruct-ERIC Centre Finland, FINStruct, and Biocenter Finland. We acknowledge the access and services provided by the Imaging Centre at the European Molecular Biology Laboratory (EMBL IC), generously supported by the Boehringer Ingelheim Foundation. We thank Jeffrey Saffitz for critical reading of the manuscript.

## Author contributions

Investigation: E.E., A.A., A.D., and J.T.H. Data Curation: E.E., A.A., A.D., and J.T.H. Methodology: E.E., A.A., V.S. Formal Analysis: E.E., E-P.K., S.K.S.T., A.A., A.D., V.S., J.T.H. Software: E.-P.K. and J.T.H. Visualization: E.E., A.A., A.D., V.S., and J.T.H. Conceptualization: E.E. and J.T.H. Supervision: E.-P.K., V.S. and J.T.H. Project Administration: E.-P.K., and J.T.H. Writing – Original Draft: E.E. and J.T.H. Writing – Review & Editing: all authors. Funding Acquisition: V.S., J.T.H.

The authors declare no competing interests. Supplementary Information is available for this paper.

Correspondence and requests for materials should be addressed to.

## Supplementary Figure legends

**Supplementary Figure 1. Micropatterning of electron microscopy grids for targeting cell–cell junctions by cryogenic electron tomography**. (**a**) A schematic representation of the micropatterning workflow. A specific pattern, placed over each grid square, is used to illuminate the grid with UV light. (**b**) Different patterns tested to guide cells to form a gap junction in the middle of the grid square. The selected pattern (a narrow hour-glass shape) is outlined in green. (**c**) A representative image of a cryo-EM grid overlaid with selected pattern. (**d**) A close-up of a cryo-EM grid after micropatterning shows hourglass-shaped pattern on each grid square. Scale bar, 100 µm. (**e**) HEK293T-Cx43-eGFP cells grown on micropatterned cryo-TEM grid. The numbers indicate the number of cells per each grid square. Scale bar, 100 µm.

**Supplementary Figure 2. Targeting gap junctions in HEK293-Cx43-eGFP cells by correlative cryogenic focused ion beam scanning electron microscopy (cryo-FIB/SEM)**. (**a**) Integrated fluorescence light microscopy image of HEK293-Cx-eGFP cells without glycerol on a cryo-EM grid prior to cryo-FIB milling. The inset shows a feature marked with an arrowhead, corresponding to a gap junction plaque. (**b**) A scanning electron microscopy (SEM) image of two cells contacting each other. (**c**) The same area after milling a 200-nm-thick lamella using a focused ion beam (FIB). (**d**) The same lamella imaged from the top. (**e**) The area indicated in *e* imaged using cryo-fluorescence super-resolution light microscopy. The signal from eGFP-tagged connexin43 is in green. (**f**) The area indicated in *f* imaged using cryogenic transmission electron microscopy (cryo-TEM). The fluorescence signal from *f* is shown overlaid on the cryo-TEM image. Interruptions in the GFP signal are indicated with white arrowheads. Scale bar, 1 µm. (**g**) A schematic diagram is shown over the same area from *g* to illustrate the areas with Cx43 channels and cavities where this signal is absent. Areas with signs of non-amorphous ice have been indicated with asterisks. (**h**) A lamella of cells with 10% glycerol imaged using cryo-fluorescence super-resolution light microscopy. (**i**) The area indicated in *h* imaged using cryo-TEM. The fluorescence signal from *h* is shown overlaid on the cryo-TEM image. Some punctate GFP signals, possibly corresponding to connexosomes (annular gap junctions), are indicated with black arrowheads. Interruptions in the GFP signal are indicated with white arrowheads. Scale bar, 1 µm.

**Supplementary Figure 3. Data processing workflow and resolution estimation**. (**a**) Data processing workflow for the reconstruction of sub-tomogram averages. (**b**) Fourier shell correlation (FSC) of the connexin cryo-ET map. (**c–d**) FSC (*c*) and an isosurface rendering (*d*) of the 80S ribosome cryo-ET map.

**Supplementary Figure 4. Cellular architecture in the vicinity of gap junctions**. (**a–f**) A 1-nm-thick slices through cryogenic electron tomography (cryo-ET) reconstructions of cell-cell junctions. Areas harbouring Cx43 channels (green), 80S ribosomes (cyan), microtubules (pink) and vesicles (yellow) have been indicated. A Golgi apparatus (G) has been labelled and outlined with a dashed line in *a*. A connexosome (annular gap junction, AGJ) in the process of being endocytosed is shown in two slices, separated by 80 nm in *e* and *f*. Scalebars, 100 nm.

**Supplementary Figure 5. Curvature in the connexin-43 lattice**. (**a–c**) Cross-sections for three different class averages of gap junction channel (GJC) lattices are shown along the axes indicated. For each, a geometrical model is shown with a surface through the centre of the lattice with the positions of the GJCs (green). The degree of curvature has been exaggerated 4× for visual clarity. (**d**) The positions of the two classes (light green and green) plotted back on one tomogram. (**e**) The same view as in *d*, showing the orientation of each particle. The particles where the blue marker is away from the viewer have a different sign of curvature compared to those where the blue marker is toward the viewer. (**f**) A close-up of *e*.

**Supplementary Figure 6. Comparison of connexin-43 and connexin-26 lattices**. (**a**) Connexin-43 (Cx43) gap junction channel (GJC) lattice, created by fitting a cryo-EM structure (PDB:7Z22) to the sub-tomogram reconstruction of the Cx43 GJC lattice patch. (**b**) Connexin-26 (Cx26) GJC lattice, created by superposing a predicted structure of the full-length Cx26 GJC on the Cx43 GJC lattice and then adjusting the channel–channel distance to the known range (77±5 Å) avoiding clashes.

**Supplementary Figure 7. Predicted structure of full length connexin-43**. (**a**) The predicted model of the full length human connexin-43 (Cx43) is shown. The regions resolved in a cryo-EM structure (PDB:7Z22) are in green. The intracellular loop (IL) and the C-terminal domain (CTD) regions present solely in the predicted structure are in grey. The phosphorylation sites are in yellow. (**b–c**) The predicted structure of the full-length human Cx43 gap junction channel is shown from the side (*b*) and from the top (*c*). The colouring is as in *a*. One helix-loop-helix (HLH) motif is indicated. The phosphorylation sites have been omitted for visual clarity. (**e**) An isosurface rendering of the intracellular stem region is shown with a transparent surface. One intracellular loop (IL) and one stalk region are indicated. (**f**) HLH motifs suggested to create the channel–channel contacts in the lateral contacts layer are shown. The local six-fold axes of symmetry are labelled with hexagons. One putative dimer of two HLHs is circled.

