## Supplementary figures and images for "In situ Structure of the Human Gap Junction"

### Supplementary Figures

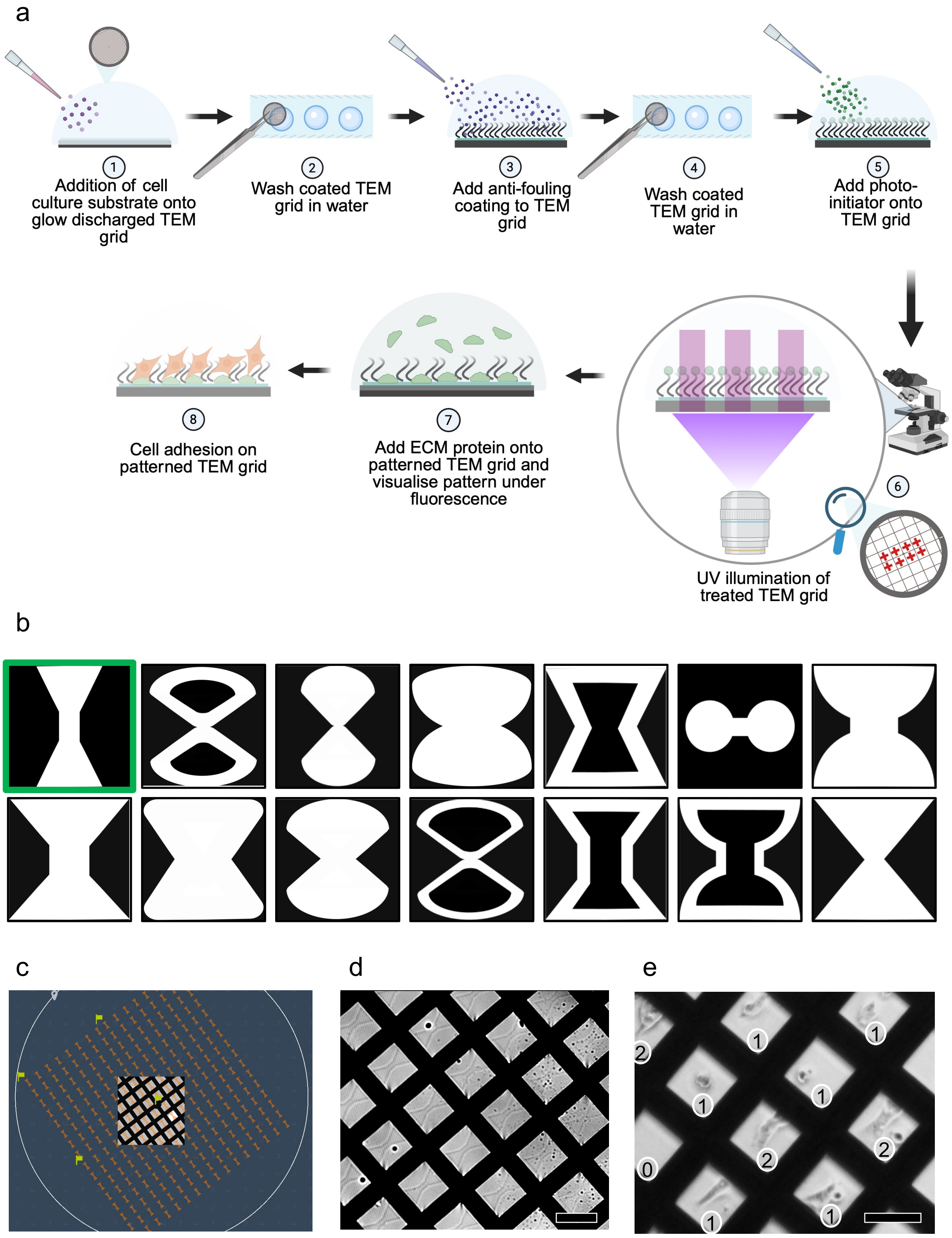

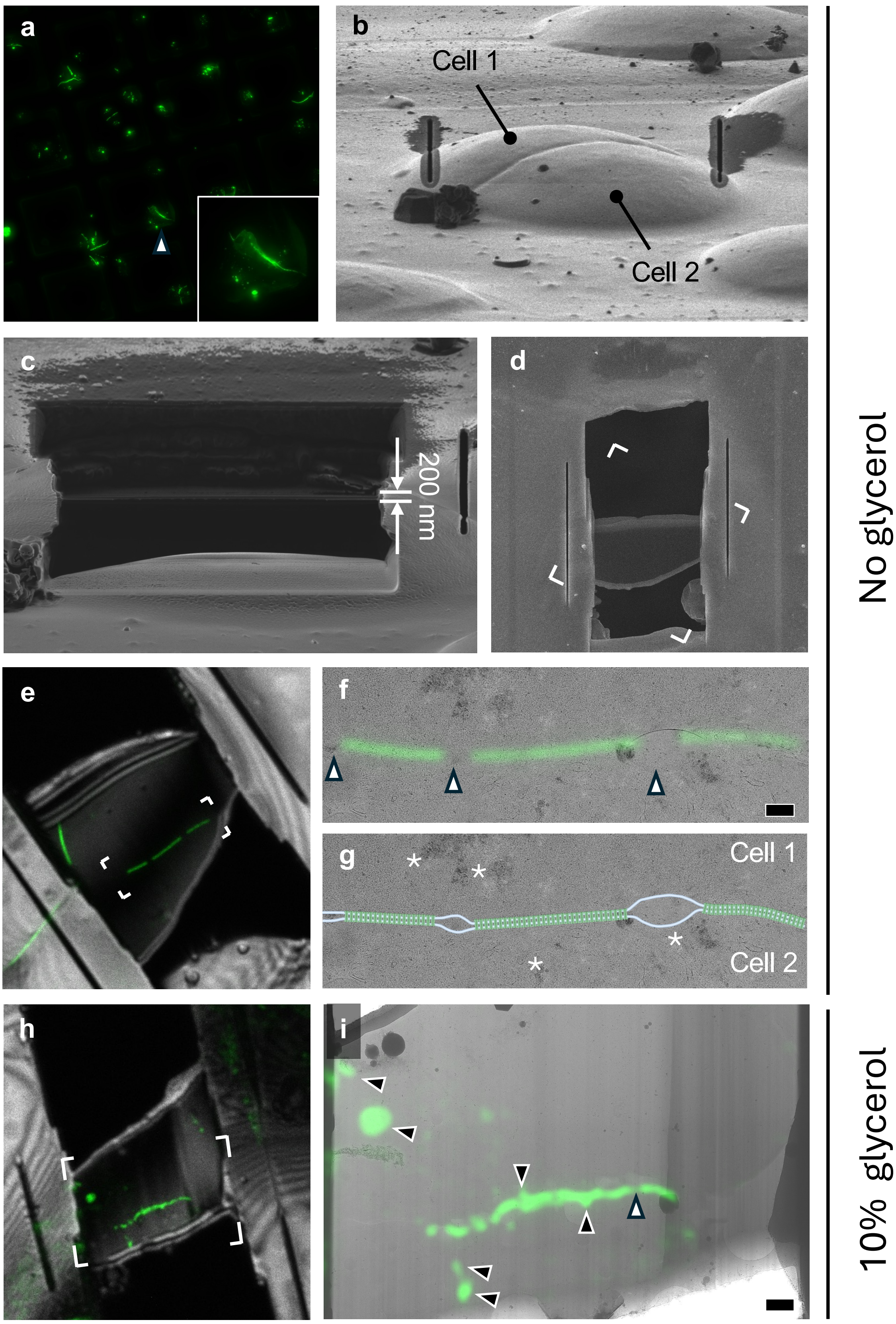

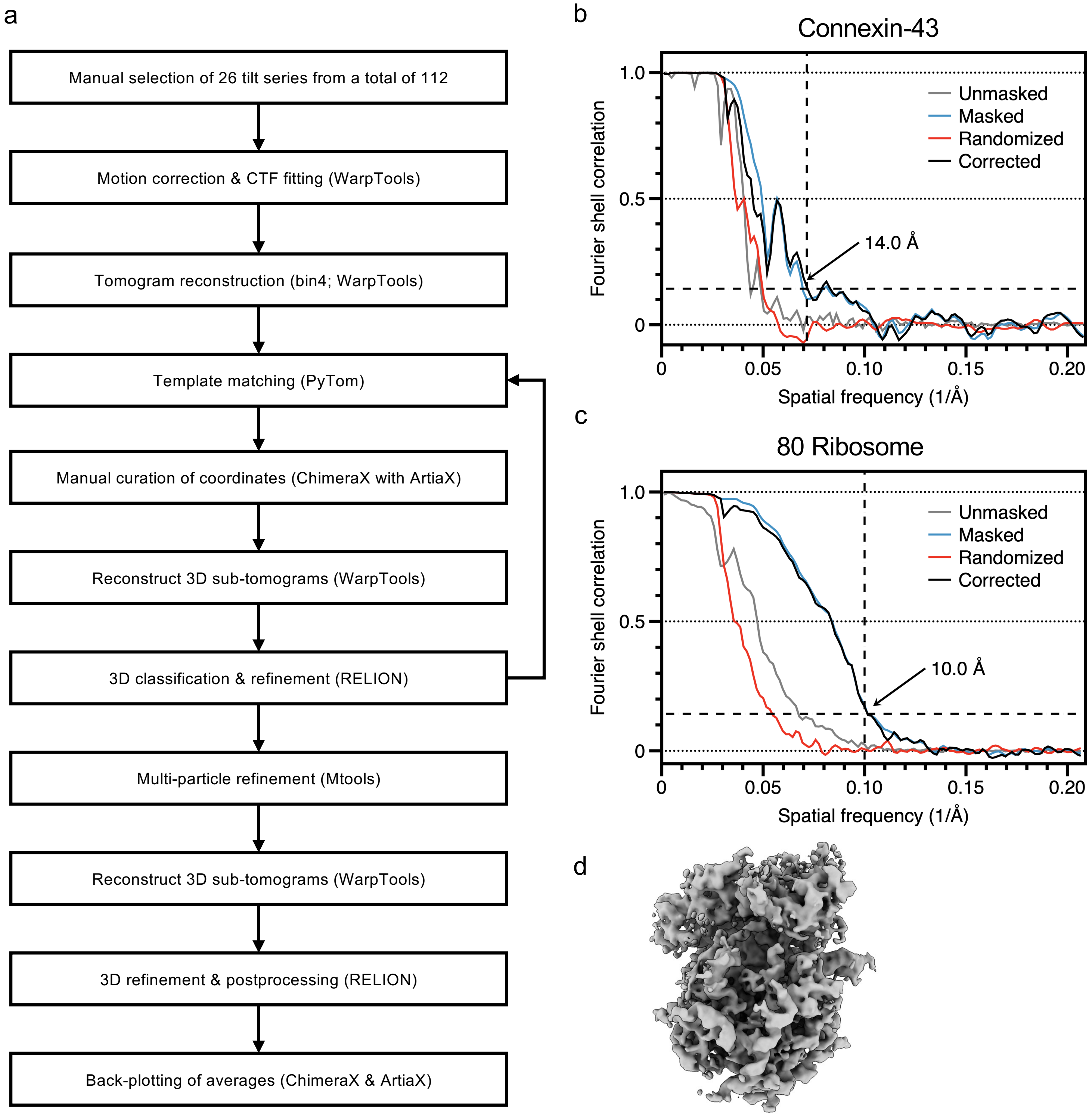

#### SFigure 4

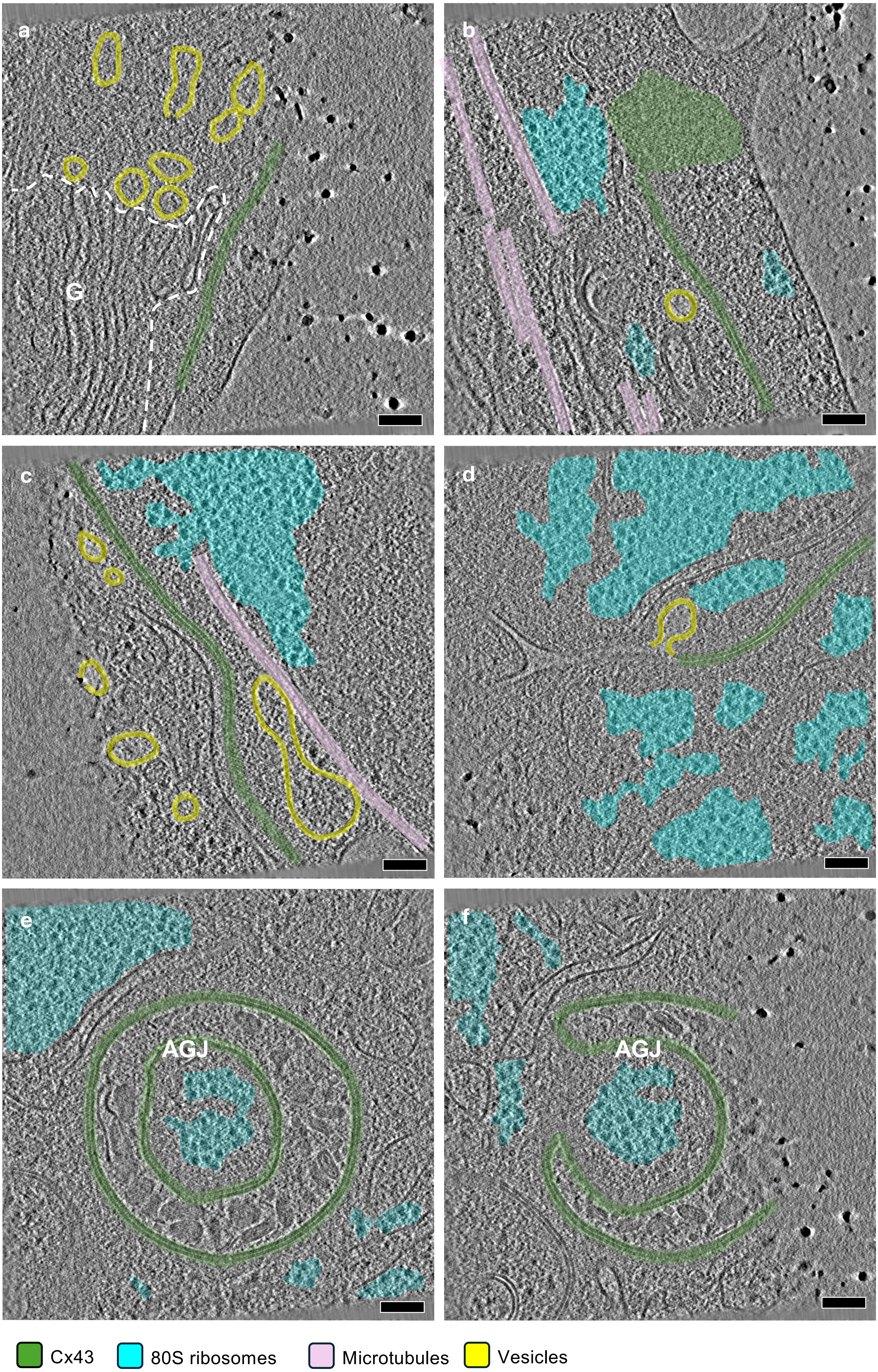

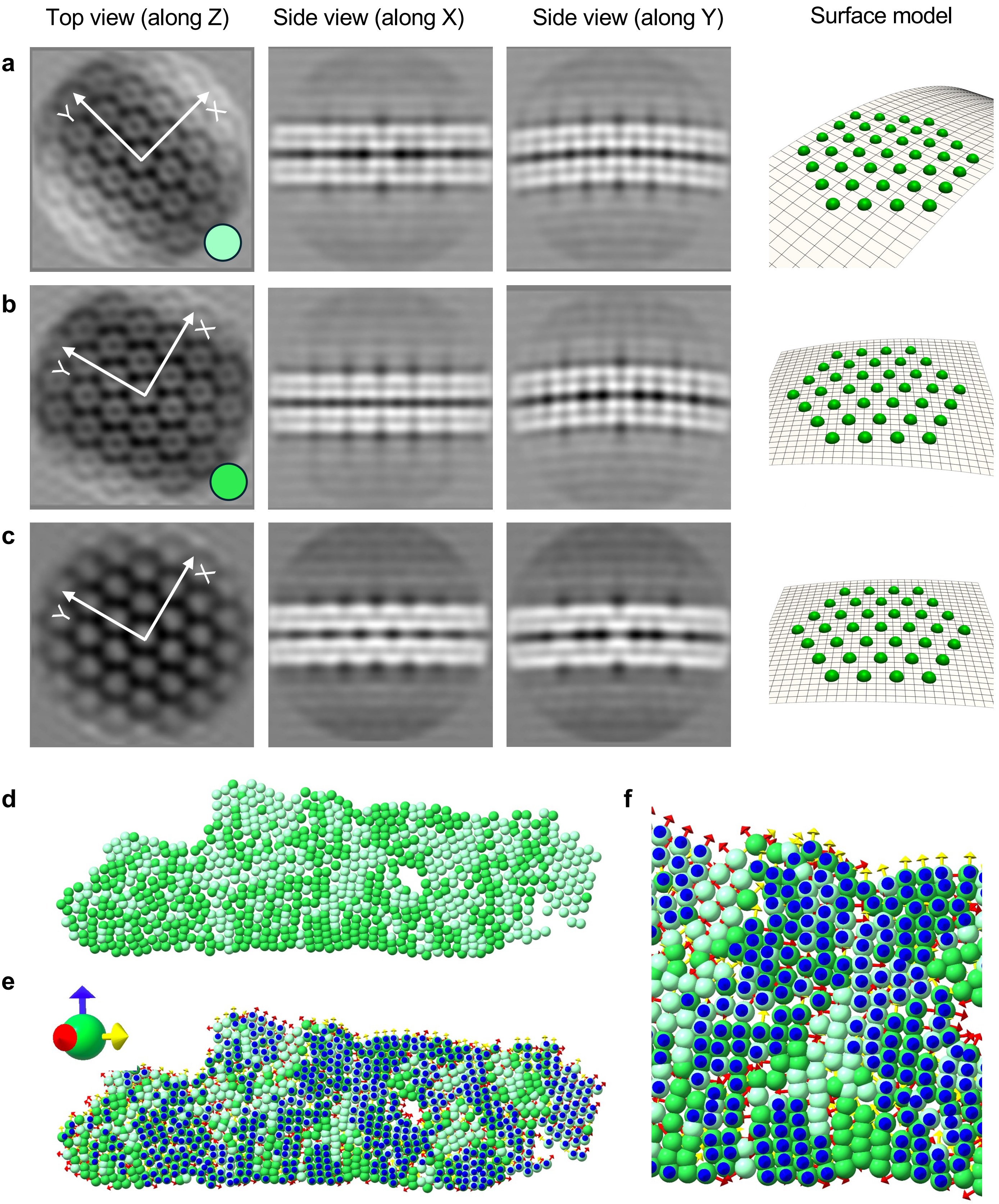

#### SFigure 6

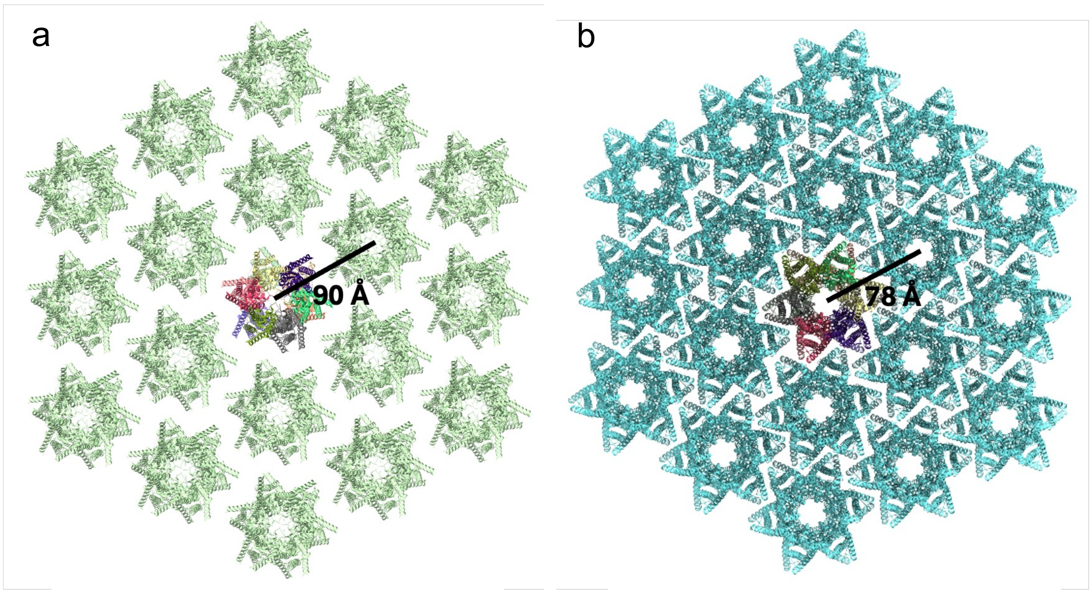


Cx43 gap junction channel lattice

Cx26 gap junction channel lattice

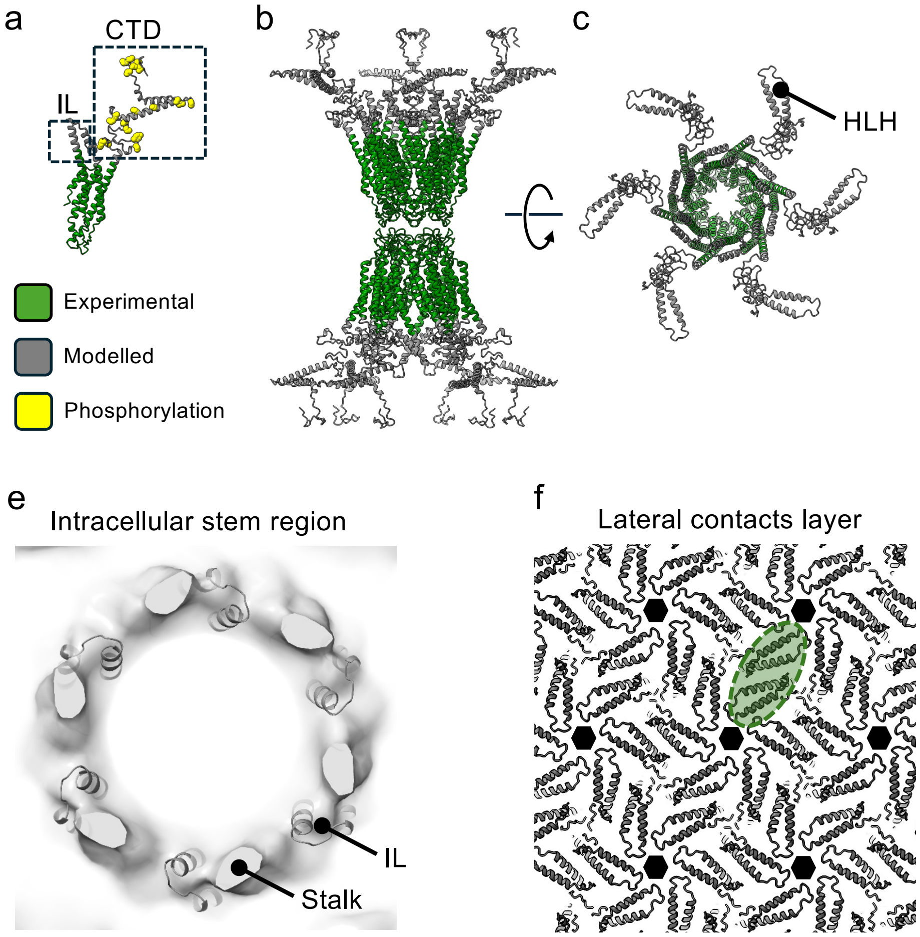
